# Functional conservation of lncRNA *JPX* despite sequence and structural divergence

**DOI:** 10.1101/686113

**Authors:** Heather M. Karner, Chiu-Ho Webb, Sarah Carmona, Yu Liu, Benjamin Lin, Micaela Erhard, Dalen Chan, Pierre Baldi, Robert C. Spitale, Sha Sun

## Abstract

Long noncoding RNAs (lncRNAs) have been identified in all eukaryotes and are most abundant in the human genome. However, the functional importance and mechanisms of action for human lncRNAs are largely unknown. Using comparative sequence, structural, and functional analyses, we characterize the evolution and molecular function of human lncRNA *JPX*. We find that human *JPX* and its mouse homolog, lncRNA *Jpx*, have deep divergence in their nucleotide sequences and RNA secondary structures. Despite such differences, both lncRNAs demonstrate robust binding to CTCF, a protein that is central to *Jpx*’s role in X chromosome inactivation. In addition, our functional rescue experiment using *Jpx*-deletion mutant cells, shows that human *JPX* can functionally complement the loss of *Jpx* in mouse embryonic stem cells. Our findings support a model for functional conservation of lncRNAs independent from sequence and structural changes. The study provides mechanistic insight into the evolution of lncRNA function.

## Introduction

Long noncoding RNAs (lncRNAs) are transcripts over 200 nucleotides in length that do not code for proteins. In contrast to protein coding transcripts, which map to only about 1.5% of the human genome, DNA sequences for lncRNA transcripts are estimated to represent 70% to 90% of the genome (Kapranov et al., 2010). Due to their low levels of expression and the general lack of functional information, lncRNAs were regarded as transcriptional noise. Only recently has the high frequency of their occurrence in the human genome and their direct relevance to various biological processes been recognized (Fatica and Bozzoni, 2014; Kopp and Mendell, 2018; Perry and Ulitsky, 2016; Rinn and Chang, 2012; Wu et al., 2017). Specifically, lncRNAs are known to be capable of scaffolding protein complexes and recruiting chromatin modifiers for transcriptional regulation (Kopp and Mendell, 2018; Mercer and Mattick, 2013; Ransohoff et al., 2017). Gene expression profiling has revealed highly tissue-specific transcription of lncRNAs and a large number of lncRNAs that are active during animal development in humans, mice, flies, and farm animals. (Derrien et al., 2012; Kern et al., 2018; Sun et al., 2013a; Wen et al., 2016). Importantly, lncRNAs have been implicated in the evolution of new genes and associated with functions in sexual reproduction (Dai et al., 2008; Gao et al., 2014; Heinen et al., 2009; Wen et al., 2016). Moreover, lncRNA functions have been shown to be conserved during embryonic development (Kapusta and Feschotte, 2014; Ulitsky et al., 2011).

LncRNAs are present in all eukaryotes; however, functions and evolution of the vast majority of lncRNA genes still remain elusive (Haerty and Ponting, 2014; Hezroni et al., 2017; Ling et al., 2015; Necsulea et al., 2014). Unlike protein-coding genes, in which functions are mostly defined by evolutionary conserved coding sequences and their flanking regulatory elements, lncRNAs are known to have poor sequence conservation (Cabili et al., 2011; Hezroni et al., 2015; Kirk et al., 2018). Hence, it has been a challenge to uncover conserved features of lncRNAs and determine underlying mechanisms for function. RNA secondary structure is one molecular feature that has recently been recognized to be important for lncRNA function (Delli Ponti et al., 2018; Fang et al., 2015; Ilik et al., 2013; Liu et al., 2017; Novikova et al., 2012; Smola et al., 2016). Yet, the direct connection between the structure and function of lncRNAs and their implications in molecular evolution remain unclear (Johnsson and Morris, 2014; Liu et al., 2017). Defining this connection has been notoriously difficult due to the lack of structural and biochemical analysis of evolutionarily related lncRNAs and the complex nature of their interactions with protein factors.

In this paper, we focus on lncRNAs involved in a mechanism of dosage compensation known as X chromosome inactivation to determine whether function is conserved and if molecular features such as RNA sequence and secondary structure influence conservation. X chromosome inactivation is the evolutionary solution to the 1X:2X dosage imbalance between XY male and XX female mammals. Outside the lineage of modern mammals, different mechanisms are used to balance the sex chromosome gene dosage. Interestingly, dosage compensation has evolved independently in divergent species and frequently uses lncRNAs as key regulators. Examples include *roX1* and *roX2* for Drosophila; *Rsx* for opossum; and *Xist*/*XIST* for mouse and human (Grant et al., 2012; Payer and Lee, 2008; Straub and Becker, 2007; Wutz et al., 2002). This suggests that RNA, unlike protein, carries functional advantages that aid and are sometimes prerequisites for biological processes such as gene dosage controls in developing embryos. Indeed, X chromosome inactivation is controlled by a genomic region known as the *X inactivation center* (*Xic*) that encodes a cluster of lncRNAs in both human and mouse genomes (Figure S1) (Brockdorff et al., 1992; Brown et al., 1992). This gene cluster has evolved from a group of protein-coding genes during the divergence of eutherians and marsupials to become the home of all lncRNAs involved in X chromosome inactivation (Casanova et al., 2016; Duret et al., 2006; Elisaphenko et al., 2008; Horvath et al., 2011). It is worthwhile to note that marsupials (e.g., opossums; Figure S1B) do not have the master regulator lncRNA *Xist* for X chromosome inactivation. In the same chromosome locus, marsupials carry the protein-coding gene *Lnx3*, which does not possess dosage compensation functions or affect the sex chromosome (Duret et al., 2006; Elisaphenko et al., 2008). However, a marsupial lncRNA, *Rsx*, was discovered to play the role of silencing the X chromosome in opossums. Although *Rsx* has no obvious sequence homology with *Xist*, the function of the two genes appears to be equivalent (Grant et al., 2012; Lee and Bartolomei, 2013; Sado and Brockdorff, 2013). Thus, the use of lncRNAs for control of mammalian X chromosome inactivation represents convergent evolution of functions that may be independent of RNA nucleotide sequences.

We took advantage of a defined molecular mechanism in mouse X chromosome inactivation which involves the direct binding of a lncRNA known as *Jpx* with a specific chromatin insulator protein, CTCF, to initiate X chromosome inactivation (Figure 1A) (Sun et al., 2013b). This model allowed us to identify the molecular features underlying the function and evolution of *Jpx*. In mice, *Jpx* has been shown to activate *Xist* (Carmona et al., 2018; Li et al., 2016; Sun et al., 2013b; Tian et al., 2010). By contrast, the function of its human homolog, *JPX*, is unknown (de Hoon et al., 2017; Migeon, 2011). As a lncRNA in humans, *JPX* is expressed in early female human embryos (Figure S2A, (Petropoulos et al., 2016)). This indicates that the gene has a role in early embryogenesis and likely functions similarly to *Jpx*. Here we will compare mouse lncRNA *Jpx* with human lncRNA *JPX* and determine their homology at the levels of nucleotide sequences, RNA secondary structures, and molecular functions. Our results indicate that despite sequence and structural divergence, the two lncRNAs function through the same biochemical mechanism.

**Figure 1.**
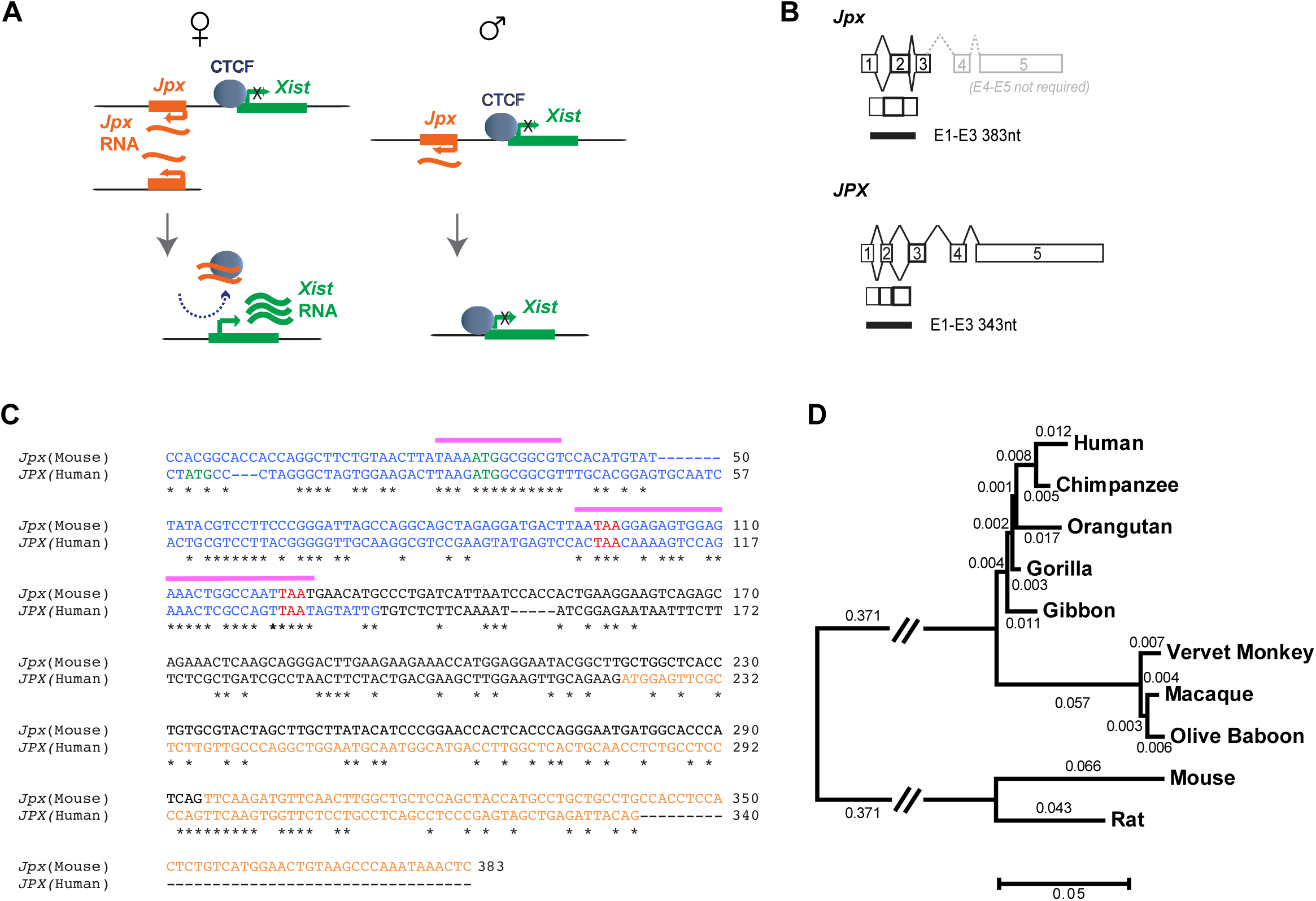
Comparative sequence analysis of mouse lncRNA *Jpx* and human lncRNA *JPX*. (A) Model of mouse lncRNA *Jpx* mechanism. LncRNA *Jpx* is transcribed upstream of *Xist*, removes CTCF from the *Xist* promoter, and activates *Xist*. This activation is dose-dependent – *Jpx* titrates away CTCF only when present in 2-fold excess, such as in female cells, and is insufficient to activate *Xist* in male cells (Sun et al., 2013b). (B) Mouse *Jpx* (Top) and human *JPX* (Bottom) gene structures and transcript isoforms. The 383nt transcript of mouse *Jpx E1-E3* is required for function (Sun et al., 2013b), which corresponds to the 343nt transcript of human *JPX E1-E3*. See also Figures S1 and S2. (C) Sequence alignment of mouse *Jpx* (Top) and human *JPX* (Bottom) transcripts; exons 1 (Blue) – exon 2 (Black) – exon 3 (Orange) analyzed by Clustal 2.1 (Larkin et al., 2007) and the alignment was manually adjusted. Asterisks mark the identical nucleotides. Pink bars label the highly conserved regions between *Jpx* and its ancestral protein homolog *UspL* (Hezroni et al., 2017). ‘ATG’ in green and ‘TAA’ in red mark potential start and stop codons, respectively. (D) Evolutionary relationships of taxa analyzed by MEGA7. Sequences were obtained from the UCSC whole genome assemblies and the evolutionary history was inferred using the Neighbor-Joining method (Saitou and Nei, 1987). The tree is drawn to scale, with branch lengths (next to the branches) in the same units as those of the evolutionary distances used to infer the phylogenetic tree. The evolutionary distances were computed using the Maximum Composite Likelihood method (Tamura et al., 2004) and are in the units of the number of base substitutions per site.

## Results

### Comparative sequence analysis suggests that human lncRNA *JPX* is functional

The molecular features and functional roles of human lncRNA *JPX* were not previously known. The GenBank annotation for human lncRNA *JPX* shows that the gene (NR_024582) contains five exons, which are similar to its mouse homolog, *Jpx* (Figure 1B) (Kolesnikov and Elisafenko, 2010; Tian et al., 2010; Tsuritani et al., 2007). Using total RNA from human ovarian cancer SKOV3iP1 cells, we were able to isolate a primary *JPX* transcript spanning exons 1-3, *JPX E1-E3*. In mice, it has been shown that the corresponding *Jpx E1-E3* is the primary isoform responsible for the function of *Jpx* in X chromosome inactivation (Lee et al., 1999; Sun et al., 2013b). More precisely, nucleotides 1-383 of the *Jpx E1-E3* sequence are necessary and sufficient for mouse lncRNA *Jpx* binding to CTCF (Sun et al., 2013b). As lncRNA *JPX E1-E3* was detected in human cells, we suspected conservation of gene structure and nucleotide sequence between *JPX* and *Jpx*. To characterize their sequence homology, we performed a pairwise sequence alignment between the critical 383nt mouse *Jpx E1-E3* with the full-length (343nt) human *JPX E1-E3* (Figure 1C). Despite an overall similarity of gene structure, including five exons in both *Jpx* and *JPX* (Figure 1B), the exact sequence identity of exons 1-3 is approximately 40%, which is much less than the average nucleotide sequence identity (85%) for protein-coding sequences between humans and mice (Gibbs et al., 2004; Makalowski et al., 1996)(one-tailed binomial test, *P* < 10^−6^). Interestingly, exon 1 of *Jpx* and *JPX* both contain remnants of protein-coding sequences similar to the chicken *UspL,* which have recently been reported as possible regulatory sequences for lncRNA function (Hezroni et al., 2017).

While a lack of sequence conservation is not surprising for noncoding genes (Necsulea and Kaessmann, 2014; Ponjavic et al., 2007; Ponting and Lunter, 2006), it is unknown how lncRNAs evolve and how noncoding nucleotide changes affect the lncRNA function. Taking advantage of fully sequenced genomes of multiple vertebrates, we searched for other homologous sequences of *Jpx E1-E3* and *JPX E1-E3* in the UCSC genome assemblies. Appropriate exon sequences for alignment were obtained from primates and murine rodents, which allowed us to look into the phylogenetic history of *Jpx*. Using the Neighbor-Joining method (Saitou and Nei, 1987), we constructed the evolutionary tree for *Jpx E1-E3* and *JPX E1-E3* in ten species (2 rodents and 8 primates), with evolutionary distances calculated from Maximum Composite Likelihood method (Tamura et al., 2004) (Figure 1D). The rate of *Jpx* sequence evolution between mouse and rat is 0.109 substitution per site, which is ∼44% lower than the neutral evolution rate of 0.196 estimated between these two rodents (Cooper et al., 2004; Gibbs et al., 2004) (one-tailed binomial test, *P* = 0.000014). This is consistent with our understanding that *Jpx* has an important functional role in mice, and thus the sequence changes have been under substantial constraint in the rodent lineages. By contrast, the evolutionary distance between rodent *Jpx* and human *JPX* is 0.796 substitution per site, which is ∼74% higher than the neutral rate of 0.457 between humans and rodents (Cooper et al., 2004) (one-tailed binomial test, *P* < 10^−6^). A more rapid nucleotide substitution between human and rodent suggests positive selection acting on the sequence of human *JPX*.

We next compared our *Jpx*/*JPX* gene tree to the recently published species tree of primates. The neutral substitution rates in the lineages leading to the hominoid (human, chimpanzee, and orangutan), as estimated from the common ancestor between hominoids and Old World Monkeys (vervet monkey, macaque, and olive baboon), are within the range of 0.026 – 0.027 (Moorjani et al., 2016). By contrast, the corresponding evolutionary rates of the *JPX* gene in the same lineages appear to be more variable, ranging from 0.009 to 0.027. Importantly in the human lineage, the nucleotide change rate is 0.012, which is two-fold higher than the neutral substitution rate of 0.0058 for humans (Moorjani et al., 2016) (one-tailed binomial test, *P* = 0.13). A larger than two-fold difference in the substitution rates is seen between the human and chimpanzee branches (bootstrap 80% over 500 replicates), which is notable given that the rates of evolution on these two lineages are estimated to be very similar with only 1.9% difference (Moorjani et al., 2016). Such observations suggest that adaptive nucleotide sequence changes have occurred in the hominoid lineages, which are supportive of a functional *JPX*, particularly in the human lineage.

Consistent with a possible role of *JPX* in regulating *XIST* within humans, re-analysis of available single-cell RNA-seq data revealed a positive correlation between *JPX* and *XIST* expression levels in human preimplantation embryos, especially in female cells of the epiblast lineage (Figure S2A; raw data obtained from (Petropoulos et al., 2016)). Such observations suggest that human lncRNA *JPX* likely functions as a positive regulator of *XIST*. Additionally, data from the Genotype-Tissue Expression (GTEx) Project showed sex-dependent expression and a positive correlation between *JPX* and *XIST* expression across 51 female samples (Pearson’s correlation *r* = 0.783, *P* < 10^−6^) (Figure S2B). Within specific tissue types, *JPX* and *XIST* activities in individual samples also showed positive correlations: *r =* 0.784 for the breast tissues (n = 290, *P* < 10^−6^); *r =* 0.690 for the pituitary tissues (n = 183, *P* < 10^−6^).

These results suggest that human lncRNA *JPX* is functionally important and that a detailed analysis of *Jpx*/*JPX* would provide a novel experimental model to understand conservation of function despite sequence diversity.

### RNA structural probing reveals divergence of *Jpx*/*JPX* homologous lncRNAs

If human lncRNA *JPX* shares similar function with its mouse homolog, it is possible that there is conservation at the RNA structural level despite a nucleotide sequence divergence. Such conservation has been supported by previous RNA structure-function studies on well-characterized noncoding RNAs, such as ribozymes and riboswitches, but thus far there has been limited analysis for lncRNAs (Ilik et al., 2013; Kirk et al., 2018; Webb et al., 2009). To explore how the sequence determines the secondary structure of RNA, we performed selective 2’-hydroxyl acylation analyzed by primer extension (SHAPE) RNA structural probing (Spitale et al., 2013; Wilkinson et al., 2006). For mouse *Jpx*, we focused on the functional lncRNA transcript *Jpx 34-347* and designed the reverse primers spanning the 314nt sequence (Figure 2A). Extension primers were also designed to probe the human lncRNA transcript *JPX 1-343* (Figure 2B). We used the SHAPE reagent, 2-methylnicotinic acid imidazolide (NAI), to modify structured *in vitro* transcribed RNA and map to residues that are accessible, such as unpaired or flexible bases (Figure 2C). As has been reported (Ilik et al., 2013; Wilkinson et al., 2006), sites of nucleotide modification can be identified as stops to primer extension by reverse transcriptase. Using radiolabeled reverse primers to generate cDNAs from NAI treated or DMSO treated RNAs, we were able to resolve the modified unpaired bases by running the reverse transcribed cDNAs through denaturing gel electrophoresis for sequencing. The intensities of the gel bands are positively correlated with the NAI modification strengths and thus reveal features of the RNA secondary structures at single-base resolution. In the SHAPE profiles for the 5’ and 3’ regions of *Jpx 34-347*, we observed segments of the base pairing in nucleotides G56-G66 and A71-C81 (Figure 2C, pex1r panel), and nucleotides C309-C317 (Figure 2C, p2r panel), suggesting possible stem-loops at these sites. Similarly for the 5’ and 3’ regions of *JPX 1-343*, we observed the base pairing in nucleotides C38-C46 (Figure 2D, pR3-4), nucleotides C276-C285 and U306-C311 (Figure 2D, pR1), which suggest corresponding stem-loop features.

**Figure 2.**
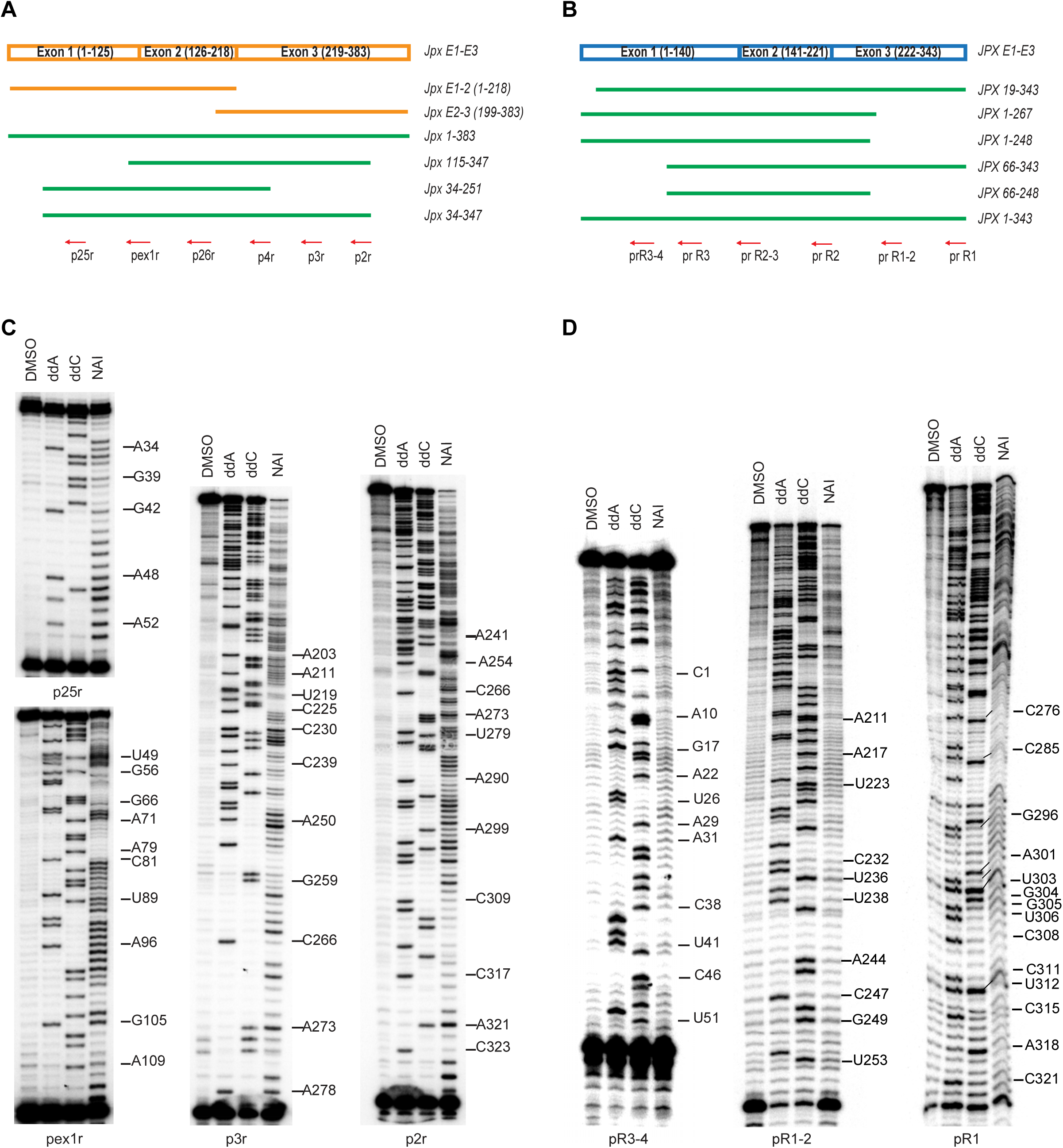
Functional domain mapping and RNA structure probing. (A) Different length *in vitro* transcribed RNA corresponding to the coverage of *Jpx*. Mouse lncRNA 220nt E1-E2 or 183nt E2-E3 (Orange) is not sufficient for protein binding *in vitro* (Sun et al., 2013b). The mouse 383nt functional *Jpx* transcript and its truncated forms (Green) are assayed with RNA EMSA. Red arrows indicate positions of the reverse primers used in SHAPE. (B) Different length *in vitro* transcribed RNA corresponding to the coverage of *JPX*. A full-length 343nt *JPX* transcript and its truncated forms (Green) are assayed with RNA EMSA. Red arrows indicate positions of the reverse primers used in SHAPE. (C) Mouse *Jpx* RNA structure probing by SHAPE: polyacrylamide gel electrophoresis (PAGE) resolves RNA footprint after treatment of RNA by either DMSO (control) or SHAPE modification reagent NAI (Spitale et al., 2013), followed by RNA reverse transcription using primers indicated and RNA hydrolysis. (D) Human *JPX* RNA structure probing by SHAPE. (C-D) At least two replicates were performed for each reaction and representative gel images are shown. Band intensity and corresponding nucleotide positions were integrated with SAFA software (Das et al., 2005; Laederach et al., 2008). SHAPE reactivities reflect single-stranded (highly reactive) and double-stranded (not reactive) states at individual nucleotides. Nucleotide labels correspond with NAI modified nucleotides.

To systemically analyze the lncRNA SHAPE profiles, we derived the SHAPE reactivity for each nucleotide after measuring band intensity, background (the DMSO lane) subtraction, and normalization (Ilik et al., 2013). By inputting all the SHAPE reactivity values into the ViennaRNA program for RNA secondary structure (Lorenz et al., 2011; Washietl et al., 2012), we were able to derive the most likely structure for *Jpx 34-347* and *JPX 1-323* based on a linear log model for pairing probabilities (Zarringhalam et al., 2012). As illustrated in Figure 3A, mouse *Jpx 34-347* RNA contains multiple stem-loops. Overall, about 50% of the nucleotides are base-paired, indicating that *Jpx 34-347* RNA is highly structured. Both the 5’ and 3’ nucleotides, 34-114 and 252-347 respectively, are involved in stem-loop formation, suggesting possible secondary structural configurations necessary for function. We asked what structural features human *JPX* may share with mouse *Jpx*. As shown in Figure 3B, *JPX 1-343* RNA indeed also contains stem-loops with more than 50% of paired bases. However, the overall structure is obviously different from the mouse *Jpx* RNA structure (Figure 3).

**Figure 3.**
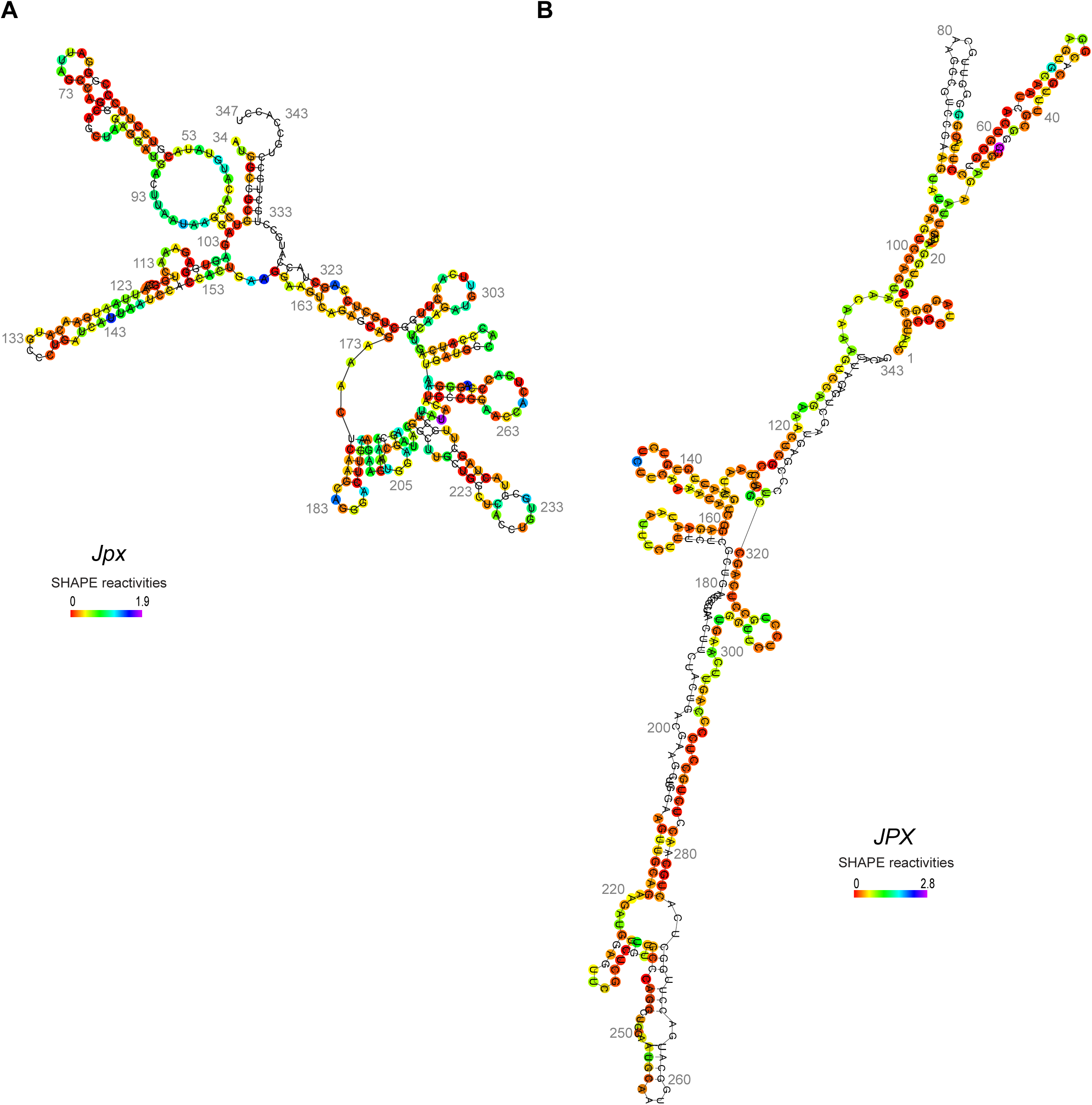
RNA secondary structures of mouse *Jpx* and human *JPX* derived from SHAPE reactivity. (A) SHAPE reactivities at individual nucleotides of mouse *Jpx* were normalized to a scale of 0 to 1.9. (B) SHAPE reactivities of human *JPX* were normalized to a scale of 0 to 2.4. (A-B) Scales were denoted with color codes at individual nucleotides. The secondary structures were drawn with the RNAprobing web server (http://rna.tbi.univie.ac.at) based on the fold algorithms in Lorenz R. et al. and Washietl S. et al. (Lorenz et al., 2011; Washietl et al., 2012) See also Figure S3.

To determine whether *in vitro* conditions may limit the RNA secondary structural probing, we performed SHAPE analysis *in vivo* with *JPX* lncRNA modified by NAI in human embryonic kidney (HEK293) cells. We note that *in vivo* RNA structural probing is particularly difficult on low abundance RNA for which results can be confounded due to the presence of more abundant RNA in the same sample (Kwok et al., 2013; Xue and Li, 2008). To enrich *JPX* for *in vivo* SHAPE profiling, we adapted a cDNA amplification step using LMPCR (ligation mediated polymerase chain reaction) which has shown to be instrumental for *in vivo* RNA structural probing (Kwok et al., 2013; Lucks et al., 2011). We chose the sequence domain *JPX* 104-172, which shares 50% nucleotide identity and corresponds to the mouse *Jpx* sequence essential for function (Sun et al., 2013b). A direct comparison between *in vitro* and *in vivo* SHAPE profiles of *JPX* 104-172 showed overlapping segments representing single-stranded RNA domains (Figure S3A). There is an overall 70% exact matching between the *in vitro* and *in vivo* RNA structural predictions (Figure S3B), with 76% (22 out of 29) single-stranded nucleotides from the *in vivo* structure falling into the loop regions predicted by *in vitro* SHAPE reactivities (Figure S3C), supporting that *in vitro* SHAPE profiling is instructive to determine RNA secondary structural features. Based on the large differences revealed by the *in vitro* SHAPE reactivities for *JPX E1-E3* and *Jpx E1-E3* (Figure 3), we conclude that human lncRNA *JPX* has diverged from mouse lncRNA *Jpx* in their overall secondary RNA structures.

### LncRNA-protein binding *in vitro* demonstrates sequence requirements for RNA function

Given the low conservation of nucleotide sequences and RNA secondary structures, we asked whether specific domains or motifs might be required for the molecular functions of *Jpx* and *JPX* lncRNAs. Utilizing the known molecular interaction between mouse *Jpx* and CTCF, we performed an *in vitro* RNA electrophoresis mobility shift assay (EMSA) and tested the binding capacity of *Jpx* RNA with regard to various truncation forms (Figure 2A). It has been reported that truncated *Jpx* RNAs, 220nt *Jpx E1-E2* or 183nt *Jpx E2-E3*, failed to bind CTCF (Sun et al., 2013b), suggesting that both halves of *Jpx E1-E3* are needed for it to function. We then removed segments of 5’ and 3’ of *Jpx E1-E3* and characterized the binding kinetics of mutant *Jpx* RNAs in comparison to the 383nt *Jpx E1-E3*. For negative control, we used a 316nt *drz-Agam-2-1* ribozyme RNA from *Anopheles gambiae* (Webb et al., 2009). As shown in Figure 4A, with increasing CTCF concentration, the full-length *Jpx 1-383* (Red) and the truncated *Jpx 34-347* (Black) both exhibited robust binding. By contrast, the truncated *Jpx 115-347* (Blue) and *Jpx 34-251* (Pink) failed to bind to CTCF. We conclude that both the 5’ sequence of nucleotides 34-114 and the 3’ sequence of nucleotides 252-347 are required for *Jpx* to bind CTCF, and that these regions may be responsible for its function.

**Figure 4.**
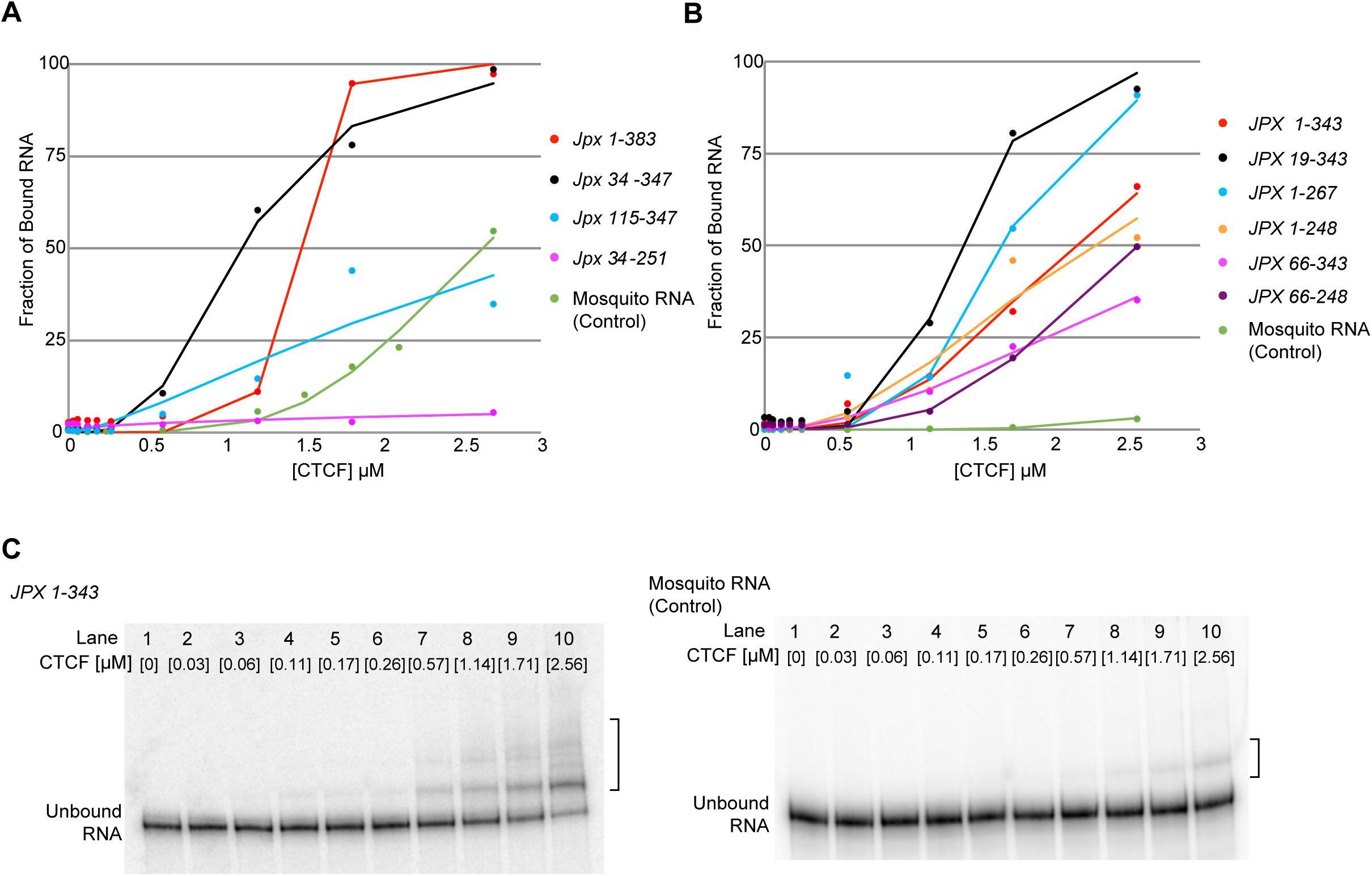
Mouse *Jpx* RNA and human *JPX* RNA are capable of binding to CTCF. (A) Binding isotherm for CTCF-*Jpx* from RNA EMSA. Binding curve was plotted as the percent bound against CTCF concentration and was fit by a nonlinear regression to a binding isotherm. The *Jpx 1-383* (Red) and the truncated *Jpx 34-347* (Black) both show robust binding, whereas 5’-truncated *Jpx 115-347* (Blue) and 3’-truncated *Jpx 34-251* (Pink) show weak or non-specific binding as compared to a 316nt control RNA (Green) from the malaria mosquito *Anopheles gambiae* (Webb et al., 2009). (B) Binding isotherm for CTCF-*JPX* from RNA EMSA. All *JPX* RNA of different lengths demonstrated favorable binding as compared to the 316nt control RNA (Green). The 5’-truncated *JPX 66-343* (Pink) and *JPX 66-248* (Purple) showed weaker binding than the 343nt *JPX* RNA (Red). (C) Representative RNA EMSA gel image detecting direct binding of *JPX* RNA and CTCF protein *in vitro*. Left panel: binding of *JPX 1-343* RNA with CTCF protein at increasing concentrations. Right panel: binding of a 316nt control RNA (from the malaria mosquito *Anopheles gambiae)* (Webb et al., 2009) with CTCF protein at increasing concentrations. RNA-protein shift is indicated by the bracket on the right side of the gel images. See also Figure S4.

We next asked whether the molecular function of human *JPX* has diverged from mouse *Jpx*, and performed EMSA on *JPX 1-343* RNA for its CTCF-binding capacity *in vitro* (Figures 4B-C and S4). We use purified recombinant CTCF with the full-length mouse CTCF protein of 736 amino acids, which is 98% identical with the human CTCF protein. In vertebrates, CTCF is highly conserved and functioning as a global transcriptional regulator in all cell types (Ohlsson et al., 2001; Phillips and Corces, 2009). Therefore, whether human *JPX* binds to CTCF the same as mouse *Jpx*-CTCF binding would inform functional importance and conservation at the molecular level. As shown in EMSA with increasing concentration of CTCF protein, *JPX 1-343* RNA was capable of binding the CTCF protein and was robustly shifted by CTCF (Figure 4C, left panel). By contrast, the 316nt control RNA of mosquito ribozyme showed weak interaction only at the highest concentration of CTCF (Figure 4C, right panel). We looked further into the binding kinetics of CTCF against various truncation forms of human lncRNA *JPX* (Figures 2B & 4B) with the goal of identifying RNA sequence domains critical for binding. As shown in Figure 4B, all *JPX* RNA truncations were able to bind CTCF in comparison to the control RNA (Green). Interestingly, the 5’ truncation form, *JPX 19-343* (Black) and 3’ truncation form, *JPX 1-267* (Blue), showed stronger binding than the full-length *JPX 1-343* (Red). Our protein-binding assays therefore indicate that human *JPX* RNA is capable of binding to CTCF, and that such a CTCF-*JPX* interaction can be robust against removal of *JPX* 5’ or 3’ RNA sequences. Overall, the EMSA results demonstrate that human *JPX* RNA has maintained, or may have even reinforced, its molecular binding capacity with CTCF protein.

Consistent with our observation of direct RNA-protein binding of *Jpx*-CTCF and *JPX*-CTCF *in vitro*, genome-wide association studies have reported CTCF-RNA interactions in both mouse and human cells (Kung et al., 2015; Saldaña-Meyer et al., 2014). Notably, mouse *Jpx* RNA was identified as one of the locus-specific interacting RNAs of CTCF in mouse embryonic stem cells by CLIP-seq (Kung et al., 2015). In addition, we also found human lncRNA *JPX* present as one of the RNA transcripts pulled-down with human CTCF by PAR-CLIP from human bone osteosarcoma U2OS cells (Supplementary data from (Saldaña-Meyer et al., 2014)). Moreover, RNA-binding regions (RBR) in CTCF have been recently reported and shown to be essential for the molecular interaction and function of CTCF in mouse and human cells (Hansen et al., 2018; Saldana-Meyer et al., 2019), which confirm CTCF binding to endogenous RNAs and the functional importance of CTCF-RNA binding in gene regulation. Our evidence of direct binding of CTCF to human *JPX* RNA similar to CTCF-*Jpx* lncRNA interaction in the mouse thus supports the functional importance and conservation of human *JPX* to its mouse homolog.

### Rescue of *Jpx*−/+ mES cell viability and morphology by a human *JPX* transgene indicates functional complementation

Since mouse lncRNA *Jpx* activates X chromosome inactivation through its binding to CTCF (Figure 1A), we therefore asked whether human lncRNA *JPX* could function equivalently for X chromosome inactivation. Using a functional complementation test, we addressed whether the *in vivo* function of human lncRNA *JPX* is equivalent to mouse lncRNA *Jpx*. We used *Jpx*−/+ female mouse embryonic stem (mES) cells for this purpose, an established *Jpx*−/+ heterozygous knockout cell line. As previously reported, *Jpx*−/+ female mES cells die during cell differentiation due to failed X chromosome inactivation, which is associated with morphology defects and a loss of *Xist* transcription (Tian et al., 2010). By introducing Tg(EF1α:JPXE1-E3) transgene expressing *JPX E1-E3* transcript into *Jpx*−/+ mutant female mES cells, we asked whether the cell viability defects caused by loss of mouse lncRNA *Jpx* could be rescued by expression of human lncRNA *JPX*. As a reference in parallel, we also introduced Tg(EF1α:JpxE1-E3) transgene expressing *Jpx E1-E3* into *Jpx*−/+ female mES cells. As shown in Figure 5A (second row), *Jpx*−/+ female ES cells receiving the vector-only transgene were dying during mES differentiation, displaying irregular and disaggregated embryonic bodies (EBs) at Day 4 (white arrows). These phenotypes were rescued by transiently transfected transgenes overexpressing *Jpx E1-E3* (third row) or *JPX E1-E3* (bottom row), with which the mutant cells formed better EBs at Day 4 and showed less dissociated cells. At Day 8 of mES differentiation, the rescue effects were most obvious. While wildtype female mES cells receiving the vector-only transgene showed EB attachment and cell outgrowth; the mutant *Jpx*−/+ had no attached EBs or cell outgrowth, and instead, had mostly disintegrated EBs and floating cells in the media. By contrast, *Jpx*−/+ female mES cells receiving either *Jpx E1-E3* or *JPX E1-E3* showed clearly attached EBs and cell outgrowth comparable to the wildtype control cells, suggesting complete reversal of cell lethality.

**Figure 5.**
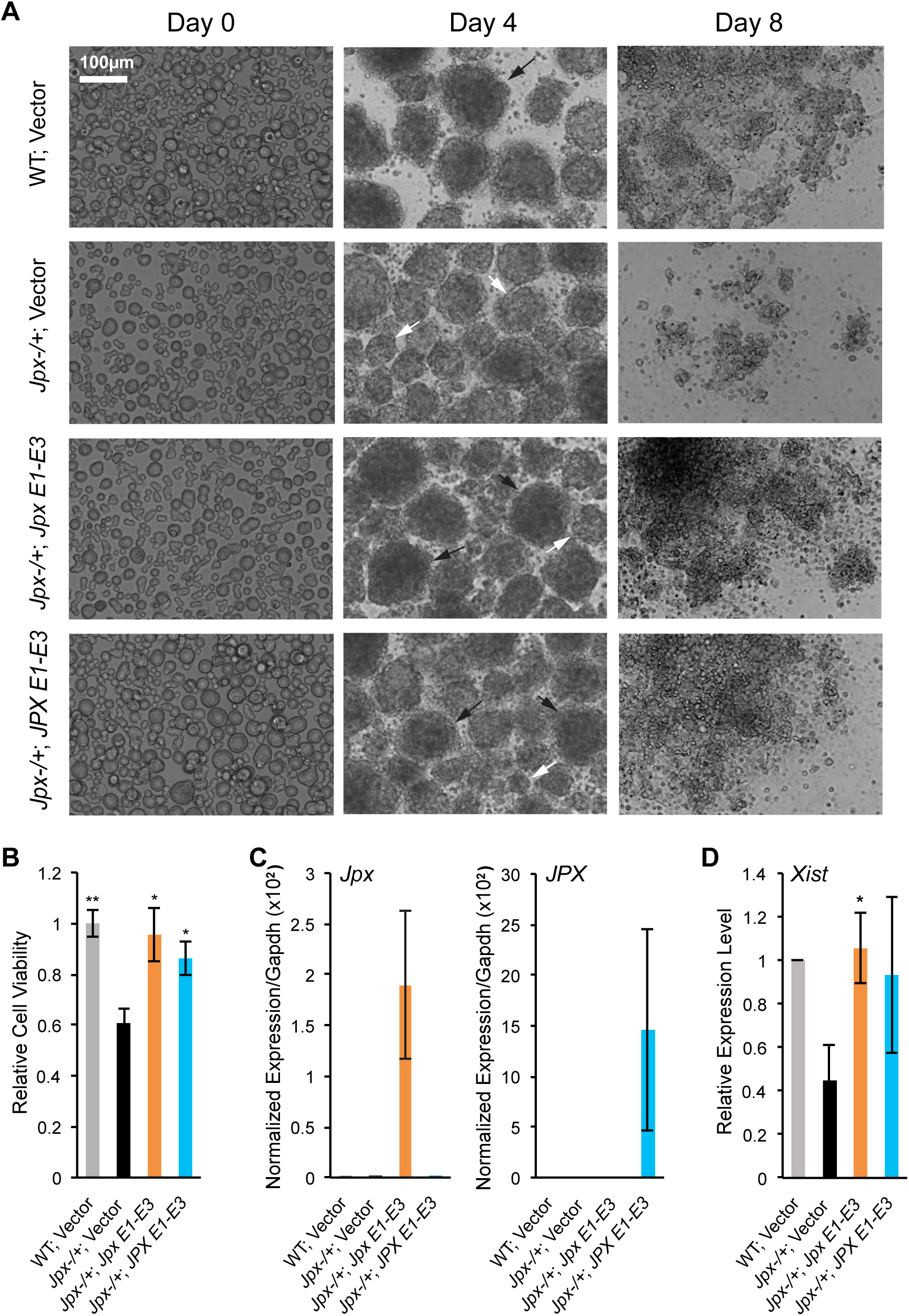
Rescue of *Jpx*-deletion in mouse ES cells by human lncRNA *JPX*. (A) Overexpression of *Jpx* and *JPX* RNA rescues outgrowth defect in *Jpx*−/+ female mutant ES cells. Wildtype (WT) control and *Jpx*−/+ female mES cells transfected with vector only, Tg(EF1α:JpxE1-E3), or Tg(EF1α:JPXE1-E3). Representative brightfield images are shown of cultures on day 0, 4, and 8 of mES differentiation. Black arrows indicate normal EBs present in cultures. White arrows indicate disintegrating EBs in the *Jpx*−/+ cultures. (B) Rescue of the cell viability defect in the *Jpx*−/+ mutant cells. At least three independent transfections were performed and average viability ± SEM are shown (*, *P* < 0.05 and **, *P* < 0.01 from one-tail paired Student *t*-tests in comparison to ‘*Jpx*−/+; Vector’). (C) Overexpression of *Jpx* and *JPX* achieved by Tg(EF1α:JpxE1-E3) and Tg(EF1α:JPXE1-E3), respectively, in the *Jpx*−/+ mutant mES cells. At least three independent transfections were performed and qRT-PCR of *Jpx* expression (left panel) and *JPX* expression (right panel) were normalized to *Gapdh* mRNA. Average expression ± SEM are shown. (D) *Xist* RNA expression rescued by *Jpx* RNA overexpression. The qRT-PCR of *Xist* expression was normalized to *Gapdh* mRNA and is shown relative to the WT level (set to “1”). Average expression ± SEM are shown from at least three independent transfections (*, *P* < 0.05 and **, *P* < 0.01 from one-tail paired Student *t*-tests in comparison to ‘*Jpx*−/+; Vector’). See also Figures S5-S7.

To validate the observed differences in EB morphology, we quantified the cell viability at Day 4. As shown in Figure 5B, mutant *Jpx*−/+ cells exhibited viability defect with ∼40% reduction as compared to wildtype female control cells of mES cell differentiation (one-tailed Student *t*-test, *P* < 0.01). In comparison, mutant *Jpx*−/+ cells receiving *Jpx E1-E3* or *JPX E1-E3* were rescued, and cell viabilities in both cases were significantly elevated (one-tailed Student *t*-test, *P* < 0.05), reaching 96% and 86% of the wildtype level, respectively. To confirm that the phenotypic rescue of *Jpx*−/+ mutant female cells was a response to the transgene expression, we assayed the *Jpx* and *JPX* RNA quantities in these transfected *Jpx*−/+ mutant female cells (Figure 5C, Day 8 shown). Mutant *Jpx*−/+ cells receiving *Jpx E1-E3* strongly expressed mouse *Jpx* but not human *JPX*, and cells that had received *JPX E1-E3* were the only ones strongly expressing human *JPX* RNA (Figure 5C). We did not observe any adverse defects when the same transgenes were expressed in wildtype female mES cells (Figure S5). EB morphology and outgrowth were normal and cell viabilities were comparable to wild type control cells carrying the empty vector (Figure S5A-B). At the molecular level, the *Jpx E1-E3* transgene in wildtype female cells induced higher *Xist* expression (Figure S5C, one-tailed Student *t*-test, *P* < 0.01), consistent with previous reports on the *trans* activation role of mouse *Jpx* on *Xist* (Carmona et al., 2018; Sun et al., 2013b). By contrast, the *JPX E1-E3* transgene did not enhance *Xist* expression in wildtype female mES cells, presumably due to the presence of intact endogenous mouse *Jpx* in these cells (Figure S5C).

A heterozygous *Jpx* deletion in the mouse female ES cell compromises overall *Jpx* expression, which leads to reduced *Xist* expression during ES differentiation (Tian et al., 2010). As shown in Figure 5D, mutant *Jpx*−/+ mES cells at differentiation Day 8 had an overall lower level of *Xist* transcripts (45% of the wildtype level). Expression of *Jpx E1-E3* transgene fully rescued *Xist* expression to 105% of the wildtype level in *Jpx*−/+ cells (one-tailed paired Student *t*-test, *P* = 0.002), consistent with the *trans* activation role of mouse *Jpx* on *Xist* (Carmona et al., 2018; Sun et al., 2013b). Expression of the human *JPX E1-E3* transgene in *Jpx*−/+ mouse cells rescued *Xist* expression to 93% of the wildtype level (one-tailed paired Student *t*-test, *P* = 0.108). This is above and beyond the confidence interval of *Xist* expression in *Jpx*−/+ female ES cells (13% – 76% of the wildtype level), indicating that human *JPX E1-E3* RNA is capable of activating *Xist* to complement the loss of mouse *Jpx* RNA in the mouse ES cells. Taken together, our results support that human lncRNA *JPX* is functionally homologous to mouse lncRNA *Jpx* in their molecular roles affecting XCI.

## Discussion

Our analyses comparing human lncRNA *JPX* with its mouse homolog *Jpx* reveal large differences in their nucleotide sequences and RNA structures but nevertheless a conservation in their molecular functions: both human *JPX* and mouse *Jpx* RNAs bind CTCF; human *JPX* can rescue *Jpx*-deletion defects in mouse embryonic stem cells. Therefore, evolutionary constraints on the sequence-structure-function linkage appear to be more relaxed and complex, consistent with current views on the genome-wide sequence evolution of noncoding RNA genes (Haerty and Ponting, 2014; Kirk et al., 2018; Necsulea and Kaessmann, 2014). A signature of positive selection acting on the human *JPX* sequence supports adaptive evolution of functional lncRNAs, which have been described in diverse organisms from *Drosophila* to mammalian species (Dai et al., 2008; Heinen et al., 2009; Kutter et al., 2012; Ponting and Lunter, 2006; Wen et al., 2016).

Early studies have reported that noncoding RNA structure can be retained across species and could possibly play a role in functional conservation, but similar analyses on lncRNAs have yet to be comprehensively performed (Ilik et al., 2013; Webb et al., 2009). To address whether structural conservation contributes into the functional conservation of *Jpx*/*JPX*, *in vitro* SHAPE analysis of both mouse lncRNA *Jpx* and human lncRNA *JPX* was performed. Upon comparison, the two structures appear divergent yet highly structured with both having stem-loop formations and ∼50% of their nucleotides base pairing. The divergence seen between these two lncRNAs may be the result of human lncRNA *JPX* evolving due to positive selection (Johnsson et al., 2014). This is supported by our protein-binding assays, which show robust interactions between CTCF and the human *JPX* RNA. A comparison of *in vivo* and *in vitro* SHAPE analyses on human *JPX* 104-172 indicated a high percentage of RNA nucleotides reactive to NAI independently of the cellular microenvironment. This region corresponds to the E1-E2 junction that is relatively more conserved in sequence and shown to bind with CTCF, thus representing a possible lncRNA domain important for function. Consistent stem-loop features obtained from *in vivo* and *in vitro* predictions suggest the structural stability of this RNA domain and support the overall structural divergence observed *in vitro* for human *JPX E1-E3* and mouse *Jpx E1-E3*.

The lncRNA-binding capability of CTCF is supported by recent characterizations of CTCF as an RNA-binding protein with specific functions in mammalian cells (Hansen et al., 2018; Kung et al., 2015; Saldana-Meyer et al., 2019; Saldaña-Meyer et al., 2014). Our analyses of *JPX/Jpx* lncRNA-CTCF binding are also consistent with the earlier report that *Jpx* RNA directly binds CTCF in activating *Xist* expression in mouse embryonic stem cells (Sun et al., 2013b). To determine whether specific RNA domains are responsible for binding to CTCF, we compare the binding affinities for the full-length *Jpx E1-E3* RNA and its mutant versions with 5’ and/or 3’ truncations, which show that both the 5’ (bases 34-114) and 3’ (bases 252-347) are necessary for CTCF-binding. Human *JPX E1-E3* RNA also binds to CTCF, and its binding capacity appears more robust against sequence deletions: the 5’ truncation mutant *JPX 19-343* and the 3’ truncation mutant *JPX 1-257* exhibit even stronger binding than the full length *JPX 1-343*. Together, our RNA EMSA results argue that CTCF-lncRNA binding with *JPX/Jpx* does not directly depend on sequence specificity.

Functional homology between human *JPX* and mouse *Jpx* is further supported by the complementary test in the mouse *Jpx*−/+ mutant ES cells. Deleting a single copy of *Jpx* gene in female mouse ES cells disrupts *Xist* upregulation and leads to cell death during ES differentiation. Exogenous expression of either mouse *Jpx* lncRNA or human *JPX* lncRNA in the mutant cells rescues *Xist* expression and cell viability. The mouse ES cell system is most suitable for the functional complementation experiment, because differentiation of mouse ES cells faithfully recapitulates the upregulation of *Xist* and the XCI process; whereas human embryonic stem cells so far are not exhibiting the establishment of random XCI or the change of XCI state during human ES cell differentiation (Khan et al., 2017; Patel et al., 2017; Sahakyan et al., 2018). Transient transfection of mouse *Jpx E1-E3* or human *JPX E1-E3* was sufficient to increase *Xist* expression and rescue the phenotypic defects of *Jpx*−/+ female cells, consistent with the *trans*-acting role of *Jpx/JPX* RNA on activating *Xist*. In contrast, transgenic mES cell lines established from stable integrations of mouse *Jpx E1-E3* or human *JPX E1-E3* transgenes show poor EB morphologies during ES cell differentiation (Figure S6). Expression of the transgenic *Jpx* or *JPX* RNA appears less efficient and variable between independent clones, which leads to insufficient rescue of cell viability in *Jpx*−/+ mutant female mES cells (Figure S7). The differences we observed between transiently transfected ES cells *vs.* stable transgenic clones likely reflect the regulatory mechanism of *Jpx* that involves the *trans*-localization of lncRNA molecules and the quantitative threshold needed for activating *Xist* (Carmona et al., 2018; Li et al., 2016).

It is interesting to note that human *JPX E1-E3* is capable of rescuing *Jpx*−/+ mES cell viability similarly as mouse *Jpx E1-E3* (Figure 5B; two-tailed paired student *t*-test, *P* = 0.469); but the efficiency of activating *Xist* is more variable with human *JPX E1-E3* than it is with mouse *Jpx E1-E3* (Figure 5D). These suggest that human *JPX* is complementary to mouse *Jpx* with regard to essential cellular functions, and that the genetic divergence may contribute to difference in the specificity in molecular interactions, which affects the regulatory efficiency on the target gene, i.e. *Xist*. This is also consistent with results from the *in vitro* protein-binding assays using CTCF.

In conclusion, through comparative sequence and functional analyses involving the homologous human *JPX*, our results have demonstrated a convergent function of *Jpx*/*JPX* between mice and humans despite a rapid divergence in the nucleotide sequences and a change of the RNA secondary structures. Our findings suggest that lncRNAs are capable of maintaining essential roles in embryogenesis and such lncRNA functions may be resistant to evolutionary constraints at both RNA sequence and structural levels.

## Experimental Procedures

### Plasmid Preparation

The Tg(EF1α:JpxE1-E3) construct was generated by PCR-cloning the *Jpx* transcript out of the cDNA library prepared from the total RNA of differentiated mES cells. The *Jpx E1-E3* isoform was cloned into pEF1/V5-His vector (Invitrogen Cat# V92020), which contains an EF-1*α* promoter for mammalian expression and a T7 promoter for *in vitro* transcription. In parallel, the Tg(EF1α:JPXE1-E3) construct was generated by PCR-cloning the *JPX* transcript out of the cDNA prepared from the total RNA of human SKOV3iP1 ovarian cancer cells. The *JPX E1-E3* isoform was cloned into the pEF1/V5-His mammalian expression vector the same way as *Jpx E1-E3*.

### RNA Electrophoretic Mobility Shift Assay (EMSA)

RNA EMSA was carried out as previously described (Cifuentes-Rojas et al., 2011; Hellman and Fried, 2007; Sun et al., 2013b) using *in vitro* transcribed RNAs uniformly labeled with ATP[α-32p] and purified recombinant CTCF protein. Specifically, *Jpx E1-E3 1-383* and truncated RNAs, or *JPX E1-E3* full-length and truncated RNAs, were *in vitro* transcribed using T7 polymerase and DNA templates PCR-amplified from Tg(EF1α:JpxE1-E3) or Tg(EF1α:JPXE1-E3) cDNA plasmid. The control RNA, a 316nt *drz-Agam-2-1* ribozyme RNA, was prepared using genomic DNA of *Anopheles gambiae* (Webb et al., 2009). Primers used are as follows.

<u>For *Jpx/JPX E1-E3* and truncated RNAs:</u>

JW_1F – TTCCCGCGAAATTAATACGACTCACTATAGGGAGATGGCGGCGTCCACATGTAT

JW_2R – AGGTGGCAGGCAGCAGGCAT

JW_4R – ATAAGCAAGCTAGTACGCAC

JW_5F – TTCCCGCGAAATTAATACGACTCACTATAGGGAGTGGCCAATTAATGAACAT

JW_21F – TTCCCGCGAAATTAATACGACTCACTATAGGGAGCCACGGCACCACCAGGCTTC

JW_22R – GAGTTTATTTGGGCTTACAG

<u>For *JPX E1-E3* full-length and truncated RNAs:</u>

hJPX-EMSA-T7+F2 – TTCCCGCGAAATTAATACGACTCACTATAGGGAGGGAAGACTTAAGATGGCGGC

hJPX-EMSA-T7+F3 – TTCCCGCGAAATTAATACGACTCACTATAGGGAGCTTACGGGGGTTGCAAG

hJPX-EMSA-R1 – CTGTAATCTCAGCTACTCGGGAG

hJPX-EMSA-R2 – GGTCATGCCATTGCATTCC

hJPX-EMSA-R3 – AGCCTGGGCAACAAGAG

hJPX-EMSA-R4 – TCGTCAGTAGAAGTTAGGCG

<u>For *drz-Agam-2-1* ribozyme RNA (control):</u>

JW_14F – TTCCCGCGAAATTAATACGACTCACTATA GCTCTGCAAATGGGGTAGGA

JW_24R – GTTTTTTCGTTTGCCGTTGAAGG

Recombinant CTCF protein was prepared and purified as previously described (Sun et al., 2013b). Mouse CTCF cDNA corresponding to the full-length 736 amino acids was cloned with C-terminal 6xHis tag into pFLAG-2 (Sigma). FLAG-CTCF-6xHis protein was induced in Rosseta-Gami B-cells (EMD Millipore) with 0.2 M of IPTG at room temperature and was purified with Ni-NTA resin (Qiagen) with 50 mM sodium phosphate (pH 8.0), 300 mM NaCl, and 250 mM imidazole. Eluates were dialyzed against 50 mM Tris-HCl (pH 7.5), 2.5 mM MgCl2, 150 mM NaCl, 0.1 mM ZnSO4, 1 mM DTT, 0.1% Tween-20, and 10% glycerol.

For gel shift (EMSA), RNAs were incubated with CTCF protein and the complexes were resolved in a 5% acrylamide gel. Gel was exposed to a phosphorimage screen (Molecular Dynamics), scanned with Typhoon phosphorimager (GE Healthcare) and analyzed with ImageQuant software (GE Healthcare). The fraction of the RNA-protein complex was plotted against the concentration of CTCF and fit with a binding equation.

### *In Vitro* and *In Vivo* RNA SHAPE (Selective 2’-Hydroxyl Acylation analyzed by Primer Extension)

RNA SHAPE was performed as previously described (Spitale et al., 2013; Wilkinson et al., 2006). Specifically, RNAs were *in vitro* transcribed from a plasmid expressing either *Jpx* or *JPX* (Tg(EF1α:JpxE1-E3) or Tg(EF1α:JPXE1-E3) respectively). Then 2pmol of purified RNA was denatured, folded, and modified with 1µmol of NAI. Immediately next was reverse transcription of the RNA and primer extension with γ-32P-ATP 5’-labeled reverse primers. Four reactions were setup for each primer extension: 1) DMSO only negative control, 2) dideoxy-ATP (ddA) in DMSO, 3) dideoxy-CTP (ddC) in DMSO, and 4) NAI in DMSO. The γ-32P-ATP end-labelled primers and ∼2pmol of RNA from the modification step were added and incubated at 95°C for the 2min annealing step followed by a 2°C/sec step-down cooling to 4°C. Reverse transcription was performed using first-strand cDNA synthesis kit containing SuperScriptIII® (2 units/µL; Invitrogen, Life Technologies).

*In vivo* SHAPE was performed as previously described with some modifications (Kwok et al., 2013; Lucks et al., 2011; Spitale et al., 2013). Briefly, HEK293T cells transiently transfected with Tg(EF1α:JPXE1-E3) for 24 hours were collected and incubated at 37°C with NAI at a 10% concentration for every 4×10^6 cells. RNA was extracted using TRIzol® (Life Technologies) according to manufacturer’s instructions. Reverse transcription was performed with non-radiolabeled reverse primers. In order to enrich the presence of *JPX*, cDNA fragments generated during primer extension with cold primers were amplified using LMPCR (ligation mediated polymerase chain reaction) (Kwok et al., 2013; Lucks et al., 2011). A linker sequence with a 5’ phosphate and 3’ 3-carbon spacer group was added to the RNA using CircLigase™ ssDNA ligase (epicentre) according to manufacturer’s instructions. PCR amplification was performed with non-radiolabeled forward primer targeting the linker sequence and γ-32P-ATP end-labeled reverse primers.

Samples were resolved on a 10% denaturing poly acrylamide SHAPE Gel (0.4mm). The gel was dried and placed into a phosphor-imaging cassette for exposure overnight, and was scanned using a Typhoon phosphorimager. Band intensities and SHAPE reactivities were calculated using SAFA software (Das et al., 2005; Laederach et al., 2008). After subtracting the DMSO background from the NAI band intensities, the average of the top 10% minus the top 2% reactivities was calculated and set to 1. This is then used to normalize the SHAPE reactivities (Ilik et al., 2013). When aligning SHAPE band positions to the transcript, the ladders (ddA and ddC) generated at the reverse transcription step are 1 nucleotide longer than the corresponding DMSO negative control and NAI samples (Wilkinson et al., 2006). RNA secondary structures are predicted with Vienna RNA Software (http://rna.tbi.univie.ac.at/cgi-bin/RNAWebSuite/RNAprobing.cgi) (Lorenz et al., 2011; Washietl et al., 2012). The extension primers used are as follows.

<u>For *Jpx*:</u>

p2r – AGGTGGCAGGCAGCAGGCAT

p3r – CTTGAACTGATGGGTGCCAT

p4r – ATAAGCAAGCTAGTACGCAC

pex1r – GGGCATGTTCATTAATTGGCCAG

p25r – TGGCTAATCCCGGGAAGGAC

p26r – CTTCAAGTCCCTGCTTGAGTTTC

<u>For *JPX*:</u>

pr R1 – CTGTAATCTCAGCTACTCGG

pr R1-2 – AGTGAGCCAAGGTCATGCCA

pr R2 – GAAGTTAGGCGATCAGCGAG

pr R2-3 – GAGACACAATACTATTAACTGGC

pr R3 – CATACTTCGGACGCCTTGCAAC

prR3-4 – CCCCGTAAGGACGCAGTGAT

LMPCR Linker Sequence – AGATCGGAAGAGCGGTTCAGCAGGAATGCCGAGACCGATCTCGTATGCCGTCTTCTGCTTG

Linker Primer F – GGAAGAGCGGTTCAGCAGGA

### Transfection of *JpxE1-E3* or *JPXE1-E3* constructs in mouse ES (mES) cells

For transient transfection, wildtype 16.7 female mES cells and the *Jpx*−/+ mutant female mES cells (Tian et al., 2010) were cultured on feeder cells in media containing LIF (leukemia inhibitory factor); the cells were collected on differentiation Day 0 (D0) and ∼1×10^6^ cells were seeded per well of a non-tissue culture treated 6-well plate for transfection. The mES cells were then transfected with 2µg of GFP plasmid (for visual confirmation of transfection) and 2.5µg of either empty vector, Tg(EF1α:JpxE1-E3), or Tg(EF1α:JPXE1-E3) plasmids per well of a 6-well plate according to the Lipofectamine® 2000 (Invitrogen) manufacturer’s instructions. Cells were cultured in feeder-free and LIF-free conditions for differentiation. On Day 1 of mES differentiation, 2mL of the transfection media is gently refreshed with regular mES media. Cells were viewed on a fluorescent microscope to assess GFP fluorescence and confirmation of successful transfection. Once transfections were deemed successful using Lipofectamine® 2000, GFP plasmid was no longer included during the transient transfection. For analyses, cells were transferred to tissue culture treated plates on D4 for EB outgrowth and allowed to grow until D8; and the cells were collected on D4 and D8 to assess viability and gene expression.

To obtain stable transgenic mES cell lines with transfection, wildtype 16.7 female mES cells and the *Jpx*−/+ mutant female mES cells (Tian et al., 2010) were seeded on feeder cells in 6-well plates at 3×10^5^ cells per well in media containing LIF. The next day cells were transfected with 2.5µg of either empty vector, Tg(EF1α:JpxE1-E3), or Tg(EF1α:JPXE1-E3) plasmids per well of a 6-well plate according to the Lipofectamine® 2000 (Invitrogen) manufacturer’s instructions while still in the stem cell state. After 24 hours, an entire well of cells were treated with 0.25% trypsin-EDTA and transferred to a 10cm dish of Neo-resistant feeder cells. Since the plasmids contain the Neomycin-resistance gene, media was switched to G418 (400ug/mL) + LIF selection media to select for cells that had stable integration of the transgenes. The medium of the cells was changed every day and after 19 days, colonies were picked and treated with 0.25% trypsin-EDTA to break up colonies and then transferred to a 24-well plate of Neo-resistant mEFs and grown out. When confluent, half of a 24-well was stocked and the other half was used for PCR screening of successful integration of plasmid. For differentiation, ∼5×10^5^ cells were seeded per well of a non-tissue culture treated 6-well plate without LIF and allowed to grow until Day 4 when they were transferred to tissue culture treated 6-well plates for EB outgrowth. Cells were collected on D4 and D8 to assess viability and qPCR analysis.

### Cell Death Assay

On Day 4 of mES differentiation, supernatant and embryoid bodies (EBs) were collected, spun down, and then broken up with 0.05% Trypsin-EDTA. Cells were then resuspended in 1mL of mES media and 20µL is taken from each sample for staining with trypan blue. Cells were counted on a Countess™ II FL Automated Cell Counter and cell viability was recorded. For Day 8 of mES differentiation, supernatant and attached cells from EB outgrowth were collected, spun down, and treated with 0.05% Trypsin-EDTA before resuspended in 1mL of mES media for cell count.

### Quantitative Real-Time PCR

Cells from one well of a 6-well plate were spun down and resuspended in 1mL TRIzol® (Life Technologies) for each respective sample. RNA was extracted and residual genomic DNA was removed with TURBO™ DNase (Ambion, Life Technologies) treatment according to manufacturer’s instructions. Reverse transcription of RNA to cDNA was performed using SuperScriptIII® (Invitrogen, Life Technologies) according to manufacturer’s instructions. Real-time PCR for *Jpx*, *JPX*, *Xist* and *Gapdh* RNA expression was performed using FS Universal SYBR Green Master (Rox) (Sigma-Aldrich) according to manufacturer’s instructions under the following conditions: 95°C for 10mins, 95°C for 15secs, 58°C for 30secs, 72°C for 30secs, then repeat steps 2 through 4 for 39 cycles. Primers used for PCR were as follows:

mJpx 76 (e1)-F – TTAGCCAGGCAGCTAGAGGA

mJpx 225 (ex2)-R – AGCCGTATTCCTCCATGGTT

hJPX E1-F – AATCACTGCGTCCTTACGGG

hJPX E3-R – GCAGGAGAACCACTTGAACT

XistNS33-F – CAGAGTAGCGAGGACTTGAAGAG

XistBP2F-R – CCCGCTGCTGAGTGTTTGATA

Gapdh-F – ATGAATACGGCTACAGCAACAGG

Gapdh-R – GAGATGCTCAGTGTTGGGGG

## Author Contributions

H.M.K., C.W., and S.S. designed research; H.M.K., C.W., Y.L., P.B., and S.S performed nucleotide sequence analyses; H.M.K. and C.W. performed and analyzed RNA structure and RNA-protein binding experiments; H.M.K. performed and analyzed mES cell experiments; D.C. and R.C.S. provided new reagents and analytic tools for RNA structure experiments; C.W., S.C., B.L., and M.E. contributed new reagents for RNA-protein binding and mES cell experiments; H.M.K, C.W., and S.S. wrote the manuscript with input from all authors.

## Acknowledgements

We thank Dr. R. Warrior for suggesting the nucleotide substitution analysis, Drs. G. MacGregor, A. Luptak, and S. Atwood for equipment, Dr. O. Razorenova for SKOV3iP1 cells, and Dr. C. Suetterlin for critique of the manuscript. H.M.K. was supported by GAANN Fellowship from the U.S. Department of Education Grant P200A120207. UCI start-up funds, Council on Research Single Investigator Foundation Grant SIIG-2014-2015-49, and the Hellman Fellowship to S.S. funded this work.

---

**Figure S1.**
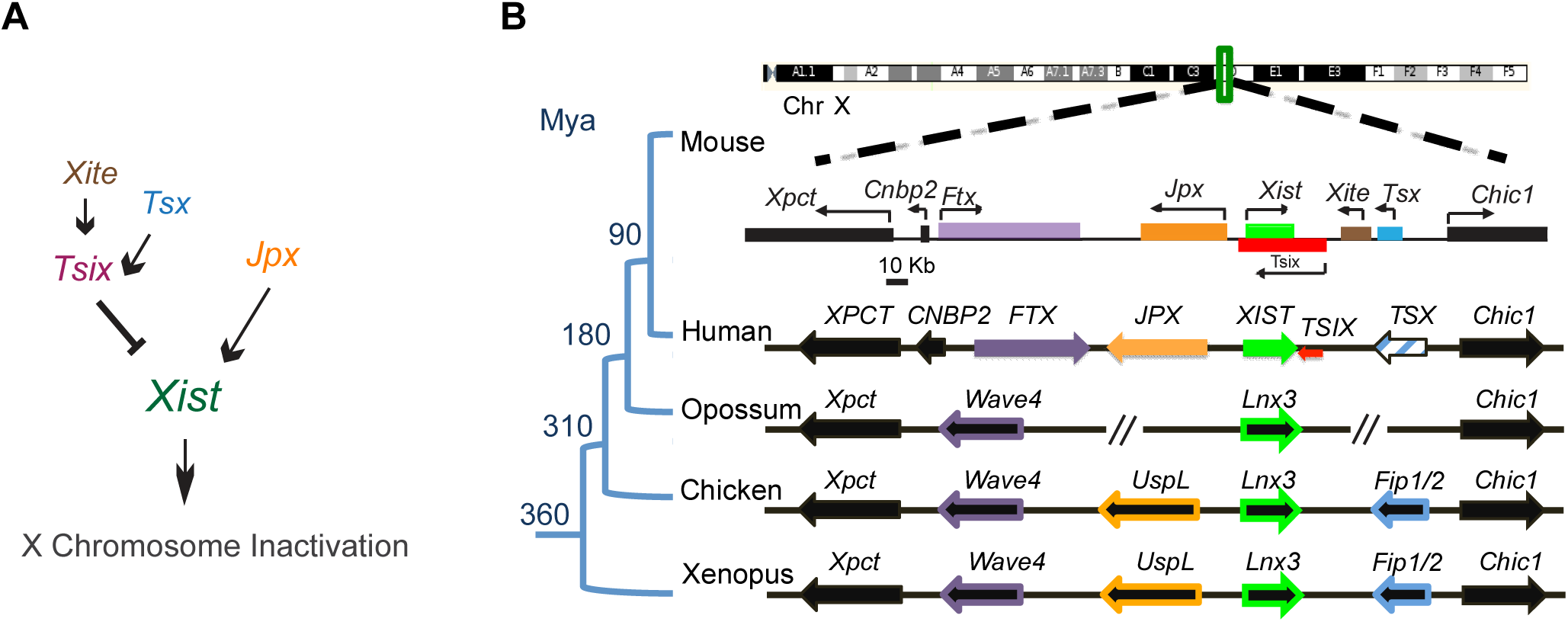
A cluster of lncRNAs at the *Xic* locus control X chromosome inactivation. Related to Figure 1. (A) *Xist* is regulated by positive and negative factors consisting of noncoding RNAs for X chromosome inactivation in the female mouse. (B) Genomic map of lncRNAs in the mouse *Xic* locus and comparison with the orthologous region in human, opossum, chicken, and frog. LncRNA genes *Ftx, Jpx, Xist, Tsix, and Tsx* are shown in solid colors with the *TSX* pseudogene in humans shown in hatched blue. Protein-coding genes *Xpct, Cnbp2, Chic1, Wave4, UspL, Lnx3, and Fip1/2* are in black with border-color matching the color of their homologous noncoding gene. Species divergence times are estimated in Mya (Million years ago), as indicated at the internal branches of the simplified phylogenetic tree. Orthologous genomes not drawn to scale; consistent with (Duret et al., 2006; Elisaphenko et al., 2008; Horvath et al., 2011).

**Figure S2.**
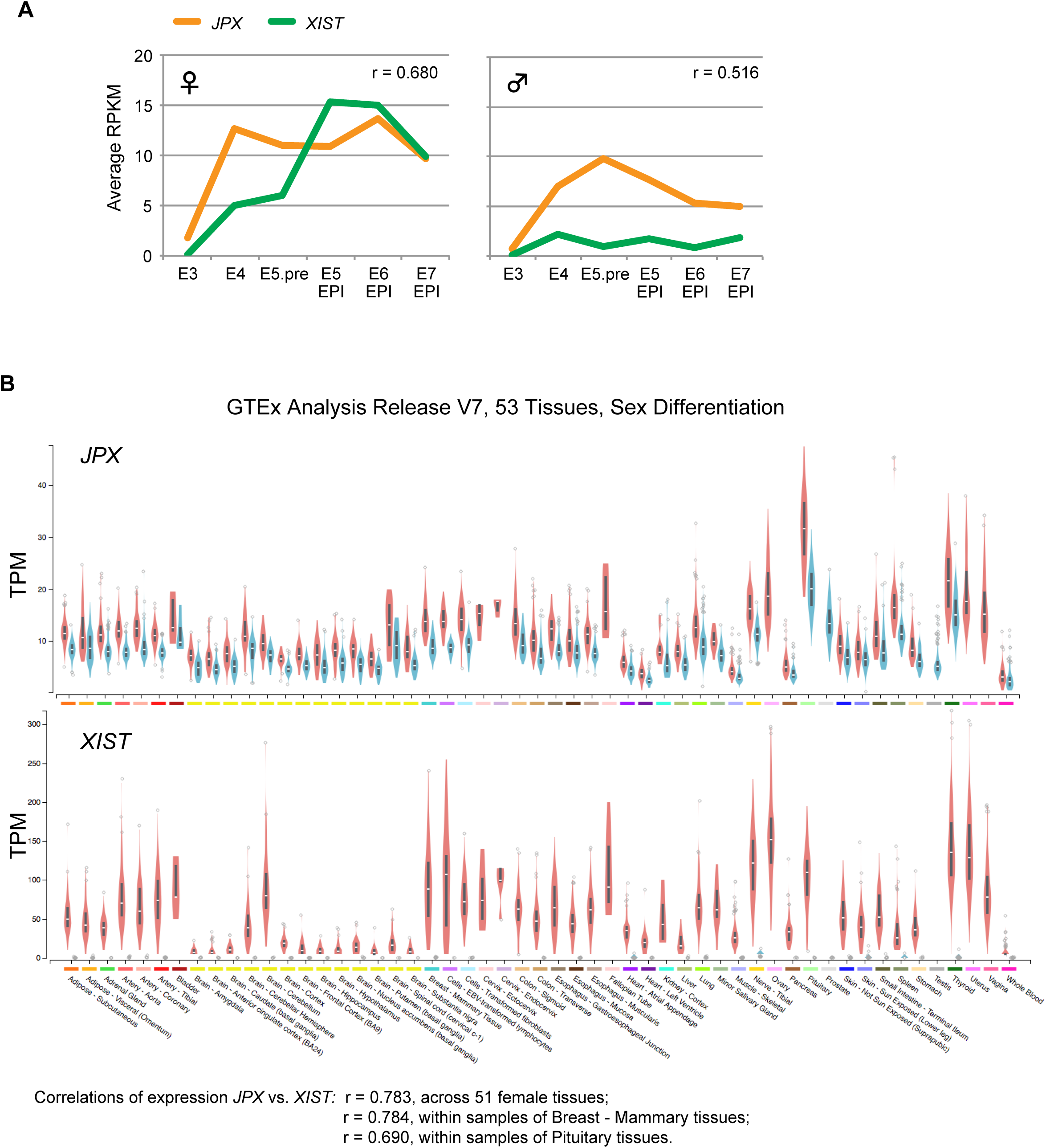
*JPX* as a possible activator of XIST in human cells. Related to Figure 1. (A) Single-cell transcript levels in human preimplantation embryos along embryonic days E3 to E7 for females (left) and males (right). Correlation of *JPX* and *XIST* expression shown as Pearson’s *r*. Analysis performed using datasets available from Petropoulos et al. (Petropoulos et al., 2016). E5.pre = Embryonic day 5 preimplantation. (B) Expression of *JPX* and *XIST* across 53 human tissues as violin plots. TPM: Transcript Per Million. Female expression is in red; male expression is in blue. Figures obtained from the Genotype-Tissue Expression (GTEx) Project, GTEx Portal (latest release version, V7).

**Figure S3.**
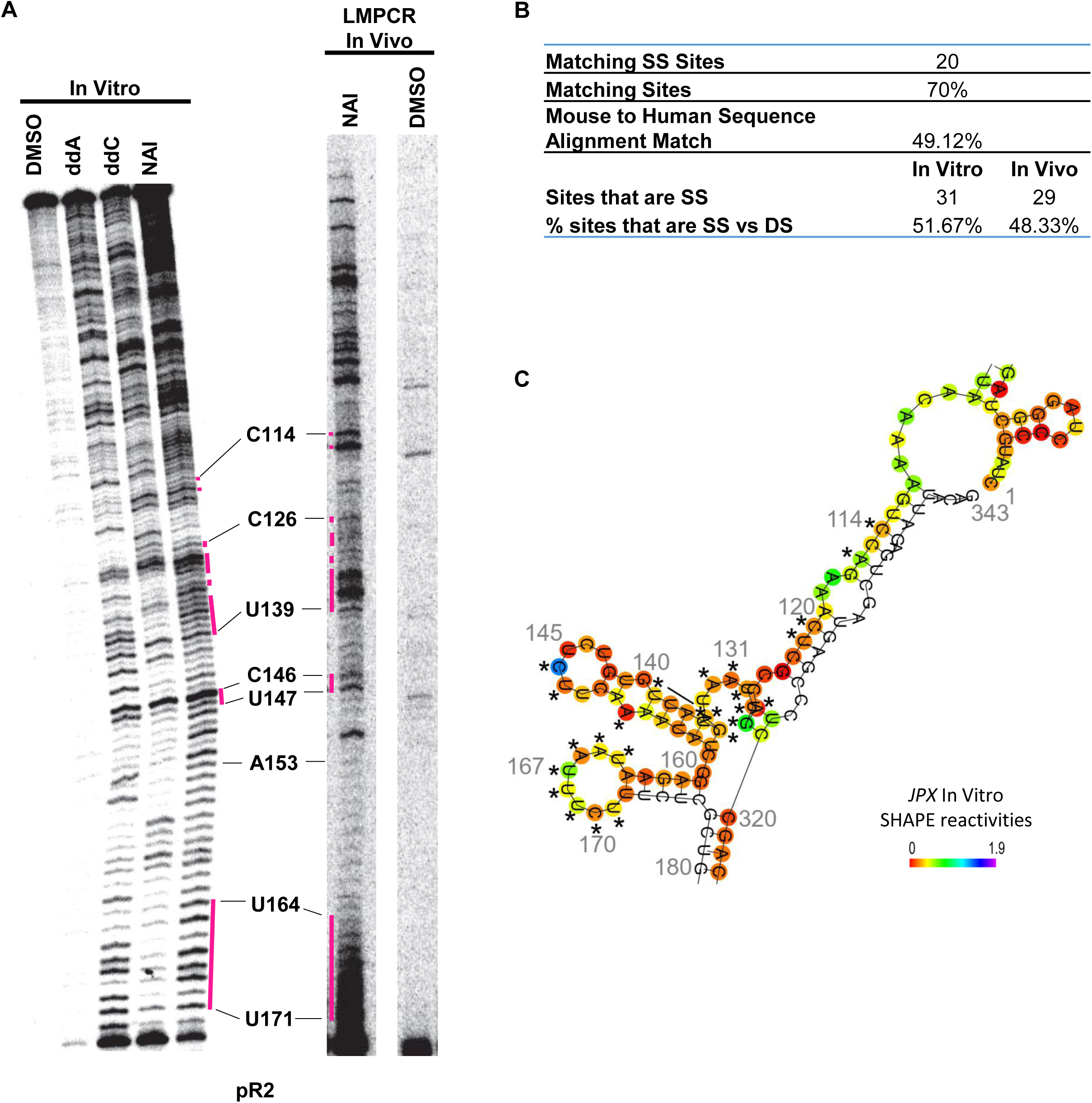
Human *JPX* RNA structure probing by SHAPE *in vitro* and *in vivo*. Related to Figure 3. (A) Polyacrylamide gel electrophoresis (PAGE) of RNA footprint after treatment of RNA by either DMSO (control) or NAI *in vitro* and *in vivo* (Kwok et al., 2013; Lucks et al., 2011; Spitale et al., 2013), followed by reverse transcription using primer pR2. Dideoxy sequencing was done using in vitro transcribed RNA. Magenta lines indicate regions of similar NAI reactive pattern between *in vitro* and *in vivo* SHAPE. These represent the single-stranded regions with unpaired nucleotides of the lncRNA *JPX*. *In vivo* SHAPE was performed using LMPCR enrichment methods (Kwok et al., 2013; Lucks et al., 2011). (B) Table of percentages of matching sites between *in vitro* and *in vivo* NAI profiles and the overall sequence conservation of the probed lncRNA region. (C) RNA secondary structure of the probed *JPX* lncRNA region as derived from *in vitro* SHAPE. *, Nucleotides reactive to NAI *in vivo* and indicated as single-stranded, as also shown in the *in vivo* SHAPE profile (A).

**Figure S4.**
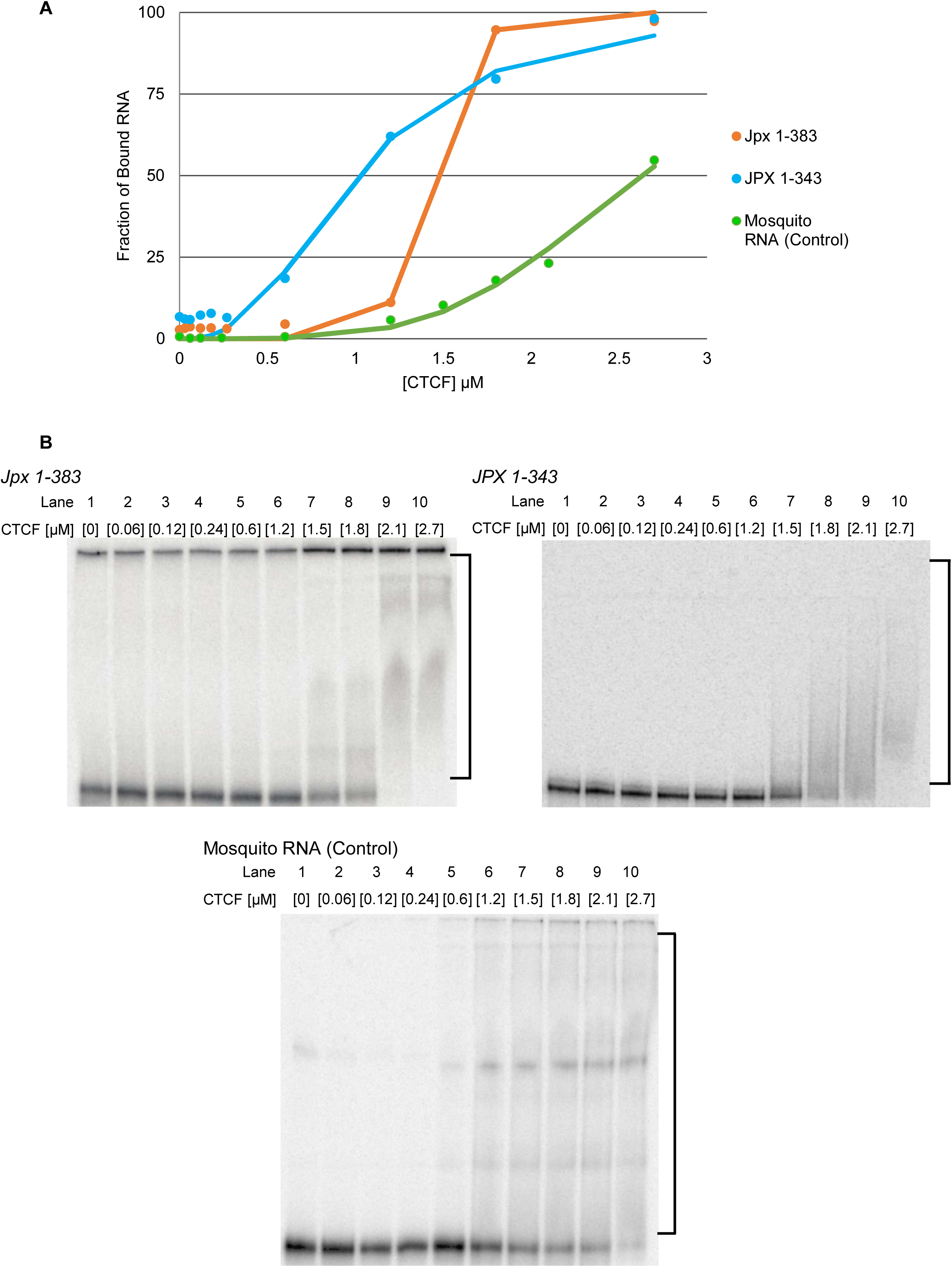
Full-length mouse and human *Jpx*/*JPX* lncRNA binding to CTCF. Related to figure 4. (A) Binding isotherm for CTCF-*Jpx*/*JPX* RNA in comparison to the control with mosquito RNA. Binding curve was plotted as the percent bound against CTCF concentration and was fit by a nonlinear regression to a binding isotherm. The *Jpx 1-383* (Orange) and *JPX 1-343* (Blue) both show robust binding as compared to 316nt control RNA (Green) from the malaria mosquito *Anopheles gambiae* (Webb et al., 2009). (B) Representative RNA EMSA gel images detecting direct binding of *Jpx*, *JPX*, and control RNA to CTCF protein *in vitro*. Top left panel: binding of *Jpx 1-383* RNA with CTCF protein at increasing concentrations. Top right panel: binding of *JPX 1-343* RNA with CTCF protein at increasing concentrations. Bottom panel: binding of a 316nt control RNA with CTCF protein at increasing concentrations. RNA-protein shift is indicated by the bracket on the right side of the gel images.

**Figure S5.**
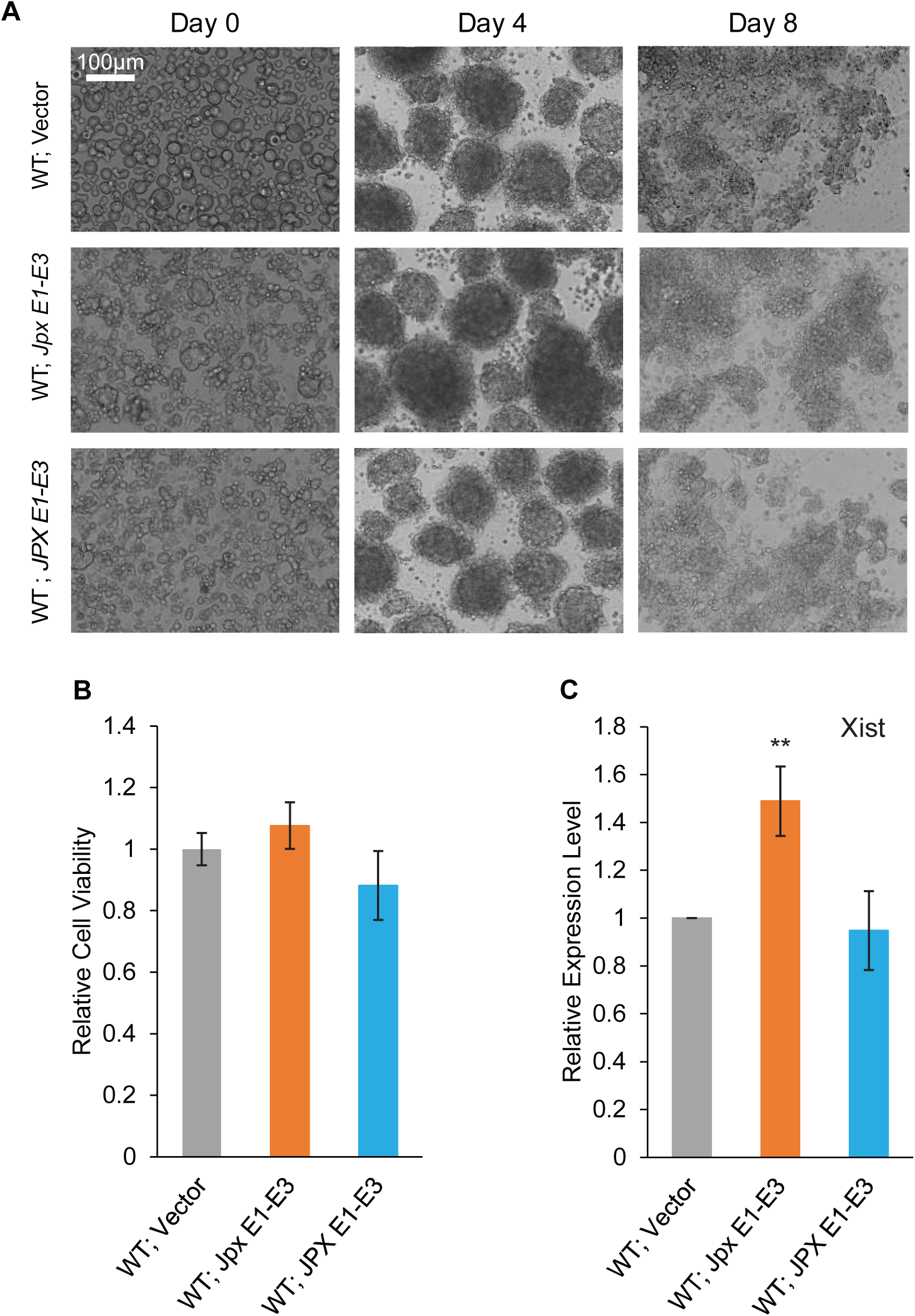
No observable defect with overexpression of *Jpx/JPX* lncRNA in wildtype mES cells. Related to Figure 5. (A) No morphological defect detected in cells overexpressing *Jpx* or *JPX* RNA. Wildtype (WT) control female mES cells transfected with vector only, Tg(EF1α:JpxE1-E3), or Tg(EF1α:JPXE1-E3). Images for WT with vector are the same ones shown in Figure 5, since the overexpression with WT cells was performed in parallel within the same experiments as the *Jpx*−/+ mutant cells. (B) WT mES cells overexpressing *Jpx* or *JPX* RNA have comparable viability with that of WT mES cells transfected with vector only. At least three independent transfections were performed and average viability ± SEM are shown. (C) No reduction of *Xist* RNA expression caused by overexpression of *Jpx* or *JPX* RNA. The qRT-PCR of *Xist* expression was normalized to *Gapdh* mRNA and is shown relative to the WT level (set to “1”). Average expression ± SEM are shown from at least three independent transfections (**, *P* < 0.01 from one-tail paired Student *t*-tests in comparison to ‘WT; Vector’). An increase of *Xist* expression by *Jpx* overexpression is consistent with previous reports (Carmona et al., 2018; Sun et al., 2013b).

**Figure S6.**
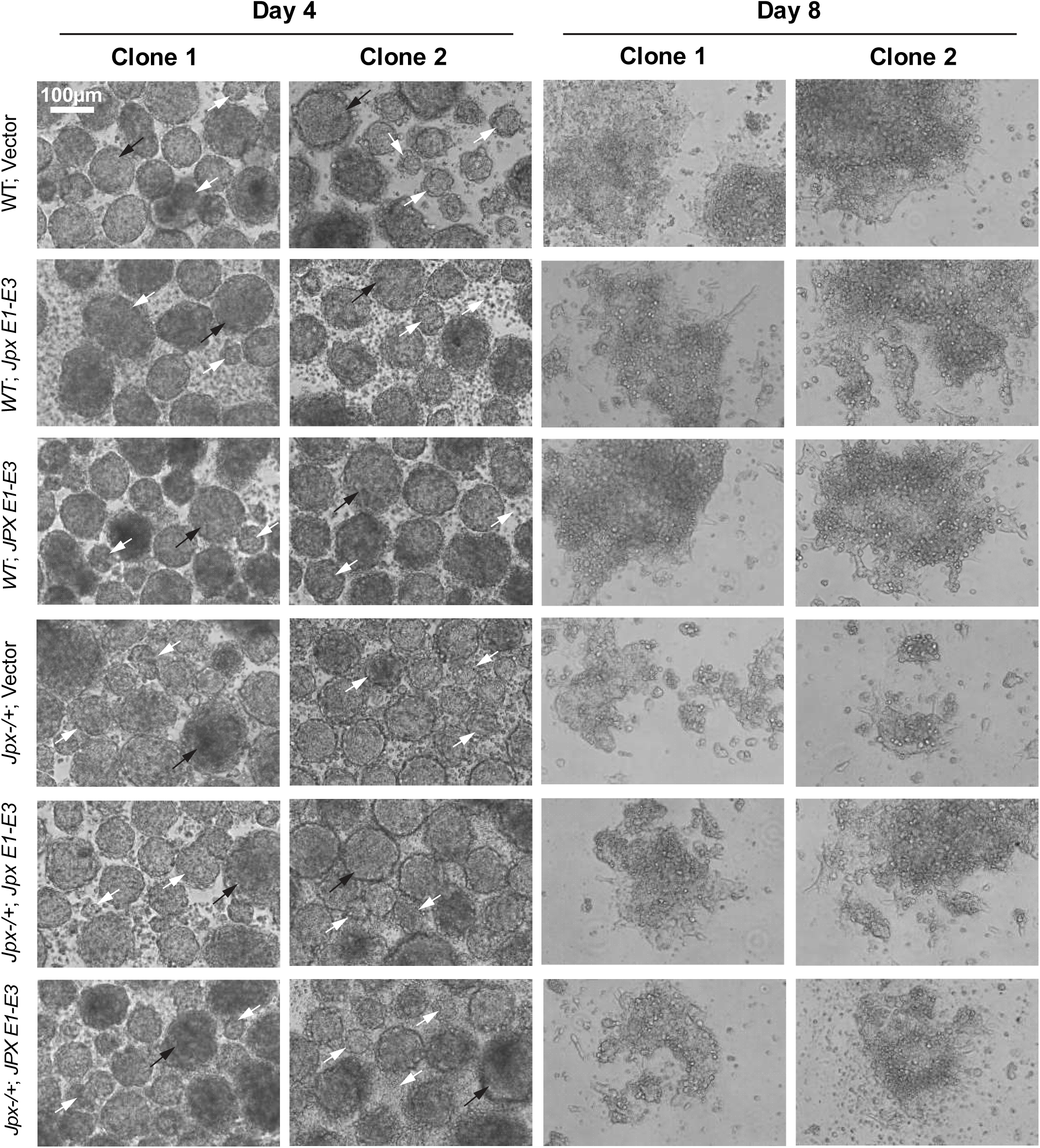
ES differentiation and EB outgrowth of stable transgenic mES cells with overexpression of *Jpx*/*JPX*. Two independent clones from each of stable transfection of *Jpx* and *JPX* transgenes in wildtype and *Jpx*−/+ female mutant ES cells. Representative brightfield images for Day 4 (EB formation) and Day 8 (EB outgrowth) of differentiated ES cells are shown for Wildtype (WT) control and *Jpx*−/+ female mES cells stably transfected with vector only, Tg(EF1α:JpxE1-E3), or Tg(EF1α:JPXE1-E3). Black arrows indicate normal EBs present in cultures. White arrows indicate disintegrating EBs in the *Jpx*−/+ cultures.

**Figure S7.**
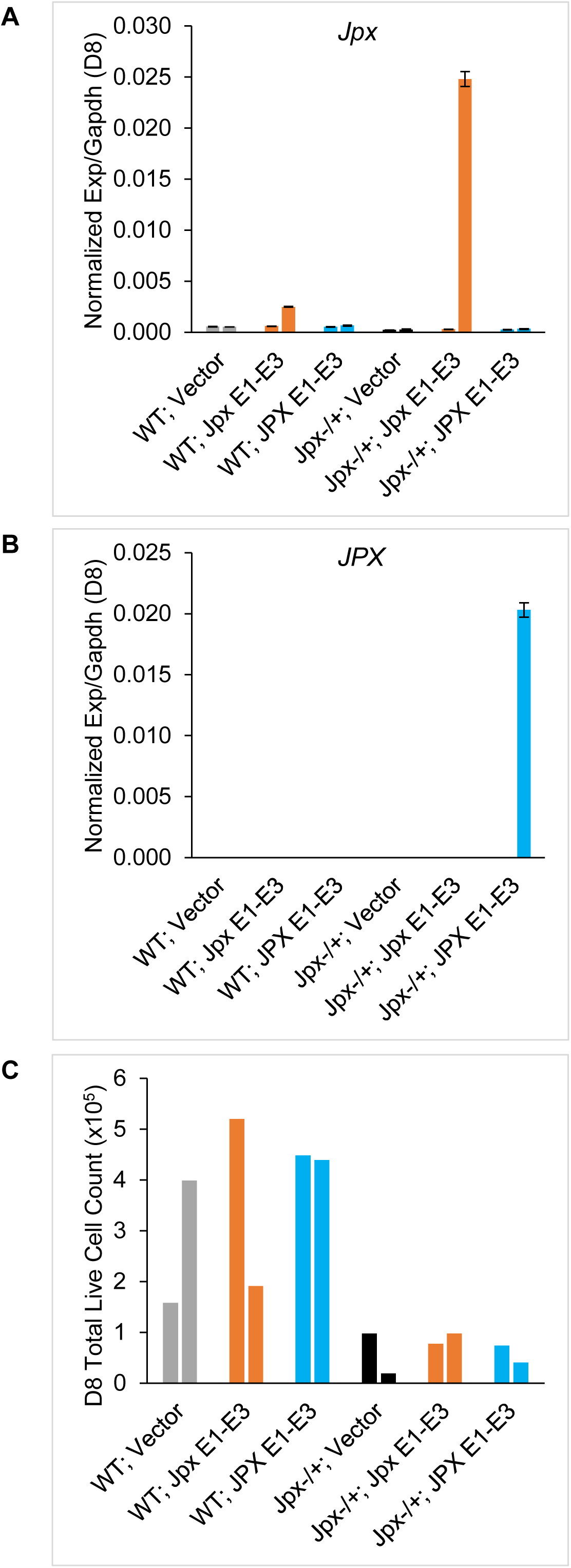
Expression of *Jpx*/*JPX* and viability of cells in transgenic mES cells with stable transfection of *Jpx*/*JPX*. Related to Figure 5. (A) Expression of *Jpx* in mES cells carrying stable transgenes. Two independent clones for each: wildtype female mES cells (WT) with vector only; WT with Tg(EF1α:JpxE1-E3); WT with Tg(EF1α:JPXE1-E3), and mutant female mES cells (*Jpx*−/+) with vector only; *Jpx*−/+ with Tg(EF1α:JpxE1-E3); *Jpx*−/+ with Tg(EF1α:JPXE1-E3). Bars represent the average of at least two qRT-PCR plate replicates of *Jpx* expression normalized to *Gapdh* for each sample. Technical replicate average expression ± SEM are shown. (B) Expression of *JPX* in the same mES cells carrying stable transgenes as in (A). Bars represent the average of at least two qRT-PCR plate replicates of *JPX* expression normalized to *Gapdh* for each. Technical replicate average expression ± SEM are shown. (C) Total live cell count on ES differentiation Day 8 for the mES cells carrying stable transgenes. Two independent clones for each type are as analyzed in (A) & (B). All samples have started with the same number of ∼5×10^5^ undifferentiated mES cells.

